# Hybridization potential between *Amaranthus tuberculatus* and *Amaranthus albus*

**DOI:** 10.1101/2021.06.11.448086

**Authors:** Brent P Murphy, Laura A Chatham, Danielle M McCormick, Patrick J Tranel

**Author notes:** Corresponding author: Patrick J Tranel 1201 W. Gregory Dr., Urbana, IL 61801.

## Abstract

The genus *Amaranthus* is composed of numerous annual herbs, several of which are primary driver weeds within annual production agricultural systems. In particular, *Amaranthus tuberculatus*, a dioecious species, is noteworthy for rapid growth rates, high fecundity, and an expanding geographic distribution. Interspecific hybridization within and between the subgenera *Amaranthus* and *Acnidia* is reported both in controlled environment and field studies, however a gap in knowledge exists with the subgenus *Albersia*. Interspecific hybridization may contribute to genetic diversity, and may contribute to the current range expansion of *A. tuberculatus*. Recently, a herbicide resistance survey of *A. tuberculatus* across five Midwestern states reported alleles of *PPX2* similar to sequences of *Amaranthus albus*, a monoecious species. Here, we seek to generate empirical data for the hybridization potential of *A. albus* and *A. tuberculatus* through replicated, controlled crosses in a greenhouse. Of 65,000 progeny screened from *A. albus* grown with *A. tuberculatus* males, three were confirmed as hybrids. Hybrids were dioecious, possessed phenotypic traits of both species, and had limited to no fertility. DNA content analysis of backcross progeny suggested a polyploid state may be required for hybrid formation. Screening of 120 progeny of *A. tuberculatus* females grown with *A. albus* identified no hybrids, though a skew to female progeny was observed. The female skew may be due to apomixis or auto-pollination, the spontaneous generation of male flowers on otherwise female plants. Our results indicate that introgression between *A. albus* and *A. tuberculatus* will occur less frequently than what has often been reported from hybridization studies with different pairs of *Amaranthus* species.

## Introduction

The genus *Amaranthus* is composed of numerous successful annual herbs of ruderal habitats. *Amaranthus* species are globally distributed, and several have naturalized throughout the world as a result of human-mediated seed flow (Sauer 1967). While much of the interest in the genus is a result of the prevalence and impact of certain species on agricultural production, they are also important components of native ecosystems (Steckel et al. 2004; Culpepper et al. 2006; Tranel et al. 2011). Within the context of annual production agricultural systems, *Amaranthus tuberculatus* and *A. palmeri* are particularly noteworthy (Tranel et al. 2011; Ward et al. 2013). These two species are both dioecious and they possess rapid growth rates, high fecundity, and expanding ranges. The factors that facilitate the expansion of these species have not been fully characterized, though range expansion is likely facilitated by herbicide resistance traits evolved within these species.

A primary selection pressure within conventional croplands is herbicides. For example, Benbrook (2016) predicted that in 2014, the total use of glyphosate within the US corresponded to 1 kg ae ha^-1^ of cropland within the country. In contrast, a typical field use rate for a single application can be considered to be 840 g ae ha^-1^, equating to over one labelled application per hectare cropland (Murphy et al. 2019). The establishment of resistant populations may both exclude native species from these habitats, and also provide a source of resistance traits to related species through hybridization. Furthermore, gene flow from these native, related species may transfer adaptive traits to the invading plants, facilitating establishment and local adaptation (Arnold 2004; Suarez-Gonzalez et al. 2018). Therefore, the adaptive potential of the gene pool, not simply a species, must be considered to predict range expansions.

Considerable genetic diversity exists within the *Amaranthus* genus, as illustrated by the naturalized range of member species. *Amaranthus retroflexus* is perhaps the most widely distributed, where the species is considered naturalized world-wide (Sauer 1967). However, most Amaranths inhabit a more narrow range. *Amaranthus tuberculatus* was historically confined to the American Midwest and Mississippi River basin (Costea et al. 2005). However, recent expansions of weedy biotypes of the species have been reported in a global and regional scale, mediated both through the direct invasion of a given biotype and the adaptive introgression of weedy traits into native, largely non-weedy genepools (Milani et al. 2020; Kreiner et al. 2019).

Hybridization across species boundaries is well documented within *Amaranthus*. A pertinent example is glyphosate resistant *Amaranthus spinosus* observed in Mississippi (Nandula et al. 2014). Hybridization with *A. palmeri* allowed the transfer and introgression of the glyphosate resistance trait into *A. spinosus*. While modern phylogenies place *A. palmeri* and *A. spinosus* as neighbors, hybridization within *Amaranthus* does not appear to be precluded even outside subgenera. Several studies have documented the hybridization potential and frequency between phylogenetically divergent *Amaranthus* species. For instance, the hybridization rate of *A. palmeri* and *A. tuberculatus*, each members of the two clades of the subgenera *Acnidia*, was observed at low frequencies both under controlled environments and field conditions (Franssen et al. 2001; Oliveira et al. 2018). In contrast, hybridization between *A. tuberculatus* and *Amaranthus hybridus*, a member of the subgenera *Amaranthus* and putative progenitor of the cultivated *Amaranthus hypochondriacus*, was observed at high frequencies under both greenhouse and field conditions (Trucco et al. 2005, 2009). Further observations of interspecific hybridization within *Amaranthus* are reviewed by Trucco and Tranel (2011).

A notable gap is observed in the case of the subgenera *Albersia*, to which the weedy species *Amaranthus albus* is a member (Stetter and Schmid 2017). No interspecific hybridization studies have been conducted with *Albersia*. However, support for gene flow between members of *Albersia* and *Acnidia* exist. A survey of resistance to herbicides that inhibit protoporphyrinogen oxidase in *A. tuberculatus* from five Midwestern states reported alleles of *PPX2* similar to sequences of *A. albus* (Nie et al. 2019). Indeed, both species are regarded as abundant weeds within the surveyed region. Plants that possessed these sequences were associated with more western sampling locations and states, where *A. albus* is expected to be more frequent. However, the frequency of interspecific hybridization between *A. albus* and *A. tuberculatus* is unknown. Here, we report on the hybridization potential of *A. albus* and *A. tuberculatus* from controlled greenhouse crosses, as measured through hybridization frequency and hybrid fecundity.

## Materials and Methods

### Population generation

The *A. tuberculatus* population PI 654437 and an in-house *A. albus* population were selected for this study. PI6554437 possesses resistance to acetolactate synthase (ALS) inhibitors, a highly heritable and selectable marker, mediated through a single amino acid substitution (W574L) in ALS (Patzoldt et al. 2005; Patzoldt and Tranel 2007). The *A. albus* population was phenotypically sensitive to imazethapyr (an ALS-inhibiting herbicide; data not shown). Reciprocal crosses were conducted between the two populations under Delnet pollen containment tents (SWM, Georgia, USA) in separate greenhouse rooms during the winter months (to limit the potential for external pollen contamination). Plants were grown under 12:12 day:night cycle, with temperature ranging from 28 to 30 C during the day and 25 to 27 C during the night in 1:1:1 soil:peat:torpedo sand mix. As *A. tuberculatus* is dioecious, pollen competition can be minimized through plant selection. However, *A. albus* is monoecious and can produce over 100,000 flowers in an indeterminate fashion; hence, seed collected from *A. albus* plants was produced under a pollen competitive environment. A total of eight *A. albus* x *A. tuberculatus* crosses were conducted across two pollination tents to obtain *A. albus* progeny, and four crosses conducted in four tents to obtain *A. tuberculatus* progeny. At maturity, plants were allowed to dry and seed obtained by manual threshing. To increase germination, seeds were surface sterilized with 50% fresh bleach, washed with deionized sterile water, and suspended in 0.1% agarose for five weeks at 4 C.

### Hybrid screening

#### Putative hybrid identification

Seeds derived from eight *A. albus* plants were screened for resistance to imazethapyr at a density of 39 viable seeds cm^-2^. Density was determined by seed weight, and the number of viable seeds was calculated based on percent germination obtained on moistened filter paper in petri dishes. Seeds were sown in growth medium (1:1:1 mixture of soil, peat, and sand) and 360 g imazethapyr ha^-1^ (Pursuit; BASF) was applied immediately after planting. Applications were made with a moving-nozzle cabinet spray chamber using an 80015 even flat fan nozzle (TeeJet Technologies), with spray volume calibrated for 187 L ha^-1^ applied 46 cm above the soil surface. After herbicide application, flats were incubated under the same greenhouse conditions described above. Surviving plants were classified as putative hybrids. Seed derived from *A. tuberculatus* female plants were classified as putative hybrids because they were produced in tents lacking male *A. tuberculatus* plants and a strong selectable marker for the *A. albus* parent was not available.

### Hybrid validation

Putative hybrids were screened with restriction fragment length polymorphisms (RFLPs) that delimit between *A. albus* and *A. tuberculatus* (Wetzel et al. 1999). The expected digest patterns for each species and their hybrids are described in Table 1. Briefly, DNA was extracted following a CTAB procedure and diluted to 50 ng uL^-1^ with spectrophotometry (NanoDrop 1000 Spectrophotometer; Thermo Fisher Scientific) (Patzoldt and Tranel 2007). RFLP assay was conducted following Wetzel et al. (1999), and imaged on a 2% agarose gel stained with GreenGlo Safe DNA Dye. Banding patterns were assessed visually to validate hybrids.

**Table 1:** *Amaranthus* species-specific digest patterns for the internal transcribed spacer region (+, restriction site present; -, restriction site absent).

| Species | DdeI | XhoI |
| --- | --- | --- |
| *A. albus* | -/- | +/+ |
| *A. tuberculatus* | +/+ | -/- |
| Hybrid | +/- | +/- |

### Backcross generations

Validated hybrids were phenotyped and backcrossed to *A. tuberculatus* PI 654437 to test hybrid fertility. BC_1_ plants were grown to the reproductive stage and tissue taken for DNA content analysis. Three floral branches per individual were harvested and stripped of floral tissue. Nuclei isolation and flow cytometry were conducted as described by Rayburn et al. (2005) with the following modifications: maize hybrid VT3 was used as an internal standard for each sample, and peak area was calculated with FCS Express software. Hybrids were phenotyped and fecundity measured through backcrosses to PI 654437. Fertility was determined through visual observation of seed production at maturity.

## Results

### Hybrids from *A. albus* as maternal parent

Eighty-six thousand seeds by weight equally derived from the eight *A. albus* parent plants were screened to identify putative hybrids. These populations possessed an average germination frequency of 75.3%, resulting in nearly 65,000 viable seed screened. A total of 13 survivors from the imazethapyr application were obtained, and screened with molecular markers. Three of these 13 survivors were identified as true hybrids (Figure 1), resulting in a hybridization frequency of 0.0046%.

**Figure 1:**
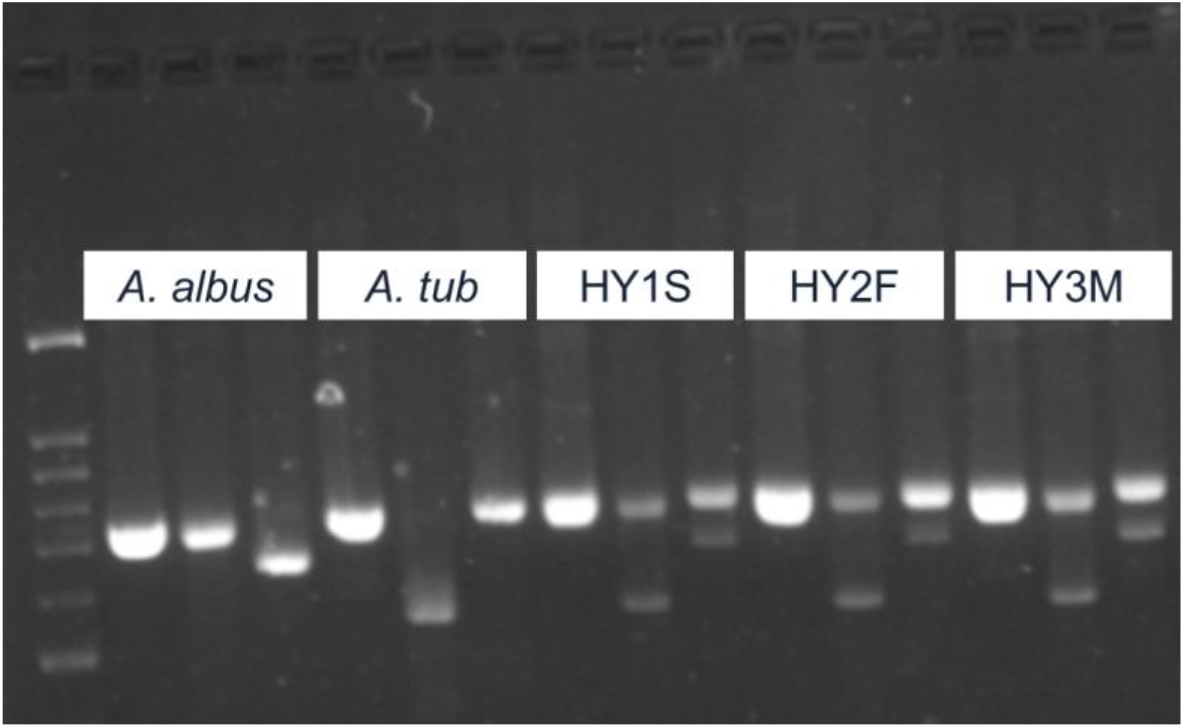
Confirmation of hybrids using restriction fragment length polymorphisms between the parental species. Band order: undigested, DdeI, XhoI.

The morphologies of mature hybrids are shown in Supplementary Figure 2. A varying degree of branching was observed in all cases, perhaps intermediate between the highly branching *A. albus* and less branching *A. tuberculatus*. In the case of hybrids HY2F and HY3M, branching was primarily observed once plants became reproductive, while HY1S began branching during vegetative stages. Stem color, which could delimit between the selected parent plants, was segregating among hybrids. HY1S and HY3M had white-green stems, similar to the *A. albus* parent, whereas HY2F had a red stem, similar to PI 654437. Limited to no fertility was observed in the case of all hybrids. All hybrids were dioecious. Of the two female plants, HY1S was fully sterile and produced no seed. HY2F was mostly sterile, though over 100 seeds were obtained from backcrosses to PI 654437. HY3M was male, and appeared to dehisce pollen, though no backcrosses were conducted due to plant staging errors.

Similar to the initial hybrids, the BC_1_ plants derived from HY2F were dioecious but otherwise possessed morphological characteristics of both parents, as shown in Supplementary Figure 3. The BC_1_ plants exhibited a higher degree of branching than is typically observed in *A. tuberculatus*. Curiously, leaves occurred throughout the terminal inflorescence. While this trait is characteristic of *A. albus*, which possesses no true terminal inflorescence, this was not observed within HY2F. All plants appeared sterile, or possessed very limited seed set (tens of seed per female). DNA content analysis revealed that all tested hybrid progeny possessed DNA contents greater than observed in either parental population (Table 2), though less than expected if HY2F was a tetraploid.

**Table 2:** DNA content of BC_1_ individuals derived from hybrid HY2F x *Amaranthus tuberculatus*.

| Sample | pg DNA 2N<sup>-1</sup> |
| --- | --- |
| <u>Parent controls</u> |  |
| *A. albus* | 1.18 |
| *A. tuberculatus* | 1.42 |
| <u>BC<sub>1</sub> individuals</u> |  |
| HY1 | 1.78 |
| HY3 | 1.85 |
| HY5 | 1.73 |

### Hybrids from *A. tuberculatus* as maternal parent

Screening of 120 seeds with the RFLP markers from the four *A. tuberculatus* female plants yielded no confirmed hybrids. Seed production of each plant was minimal, with yields ranging from tens to low hundreds of seeds produced, whereas tens to hundreds of thousands of seeds are expected when pollinated by an *A. tuberculatus* male. Progeny were grown to maturity and gender ratios calculated through visual assessment. Only 16 of the 120 plants were males, whereas progeny from PI 6354437 exposed to *A. tuberculatus* pollen were equally divided between males and females (Table 3).

**Table 3:** Gender ratios of progeny from *Amaranthus tuberculatus* females grown in the presence of *Amaranthus albus* as a pollen source.

| Female | Progeny |  | P-value<sup>a</sup> |
| --- | --- | --- | --- |
|  | Male | Female |  |
| ACR10 | 1 | 5 | 0.102 |
| ACR20 | 9 | 36 | <0.001 |
| ACR3-5 | 2 | 21 | <0.001 |
| ACR3-1 | 4 | 42 | <0.001 |
| Control<sup>b</sup> | 29 | 28 | 0.894 |

<sup>a</sup>Chi-square test was conducted against an expected 1:1 male:female ratio.
<sup>b</sup>Random subset of progeny of an *A. tuberculatus* x *A. tuberculatus* cross.

## Discussion

Three hybrids between *A. albus* and *A. tuberculatus* were successfully identified. The rate of hybridization (0.0046%) observed within this study is markedly lower than that reported from other interspecific crosses conducted within the genus. Crosses conducted between representatives of the two clades of the subgenus *Acnidia, A. palmeri* and *A. tuberculatus*, resulted in hybridization rates of 1% (Franssen et al. 2001). Crosses of *A. tuberculatus* outside of *Acnidia*, to *A. hybridus* of the *Amaranthus* subgenus, resulted in hybridization rates of 5% (Trucco et al. 2005). Interestingly, molecular phylogenies place *A. albus* as being more closely related to *A. tuberculatus* than to *A. palmeri* (Stetter and Schmid 2017), and *A. albus* has a matching chromosome number to *A. tuberculatus* (2N = 32), while *A. palmeri* does not (2N = 34) (Grant 1959). Nevertheless, major fertility issues were observed both within the initial hybrids as well as in the BC_1_ population. These results suggest that, while the cross can happen, *A. tuberculatus* is not within the primary gene pool of *A. albus*. Furthermore, flow cytology suggests that elevated DNA content was observed within the BC_1_ population. Perhaps a polyploid state is necessary to produce hybrids between *A. albus* and *A. tuberculatus*. Under the hypothesis that chromosome doubling of at least one set of gametes is required, the rate of hybridization would be the probability that the gamete is doubled, and the probability that *A. tuberculatus* will outcompete *A. albus* pollen, resulting in reduced hybridization frequencies.

Hybrid plants possessed features of both species, though were most similar to *A. tuberculatus*. Dioecy appears dominant, which is consistent with other interspecific crosses within the genus (Trucco et al. 2005). Apical dominance was observed in all hybrids during the vegetative stage, though this dominance weakened during the reproductive stage. As such, apical dominance appears dominant over the extensive lateral branching pattern of *A. albus*, which results in its “tumbleweed” morphology. Stem color was variable amongst hybrids, an indication that traits from both parents were expressed within the hybrid plants.

While theoretically possible, the low frequency of hybridization and the low frequency of fecundity across multiple generations suggests that *A. albus* is unlikely to contribute adaptive traits towards the expansion of *A. tuberculatus*. However, herbicide resistance traits could provide a qualitative fitness advantage to overcome these boundaries within agricultural systems. There is no reported case of herbicide resistance in *A. albus* within the continental US (Heap 2020). The origin of the ‘tumble-type’ *PPX2* allele observed by Nie et al. (2019) remains unresolved, but could indicate that even extremely low rates of hybridization and low viability of hybrids is still sufficient to allow gene introgression between these two species. Of the member species of *Albersia*, only *A. albus* and *Amaranthus blitoides* are considered agronomically important weeds in American Midwest, though neither is noted for herbicide resistance. An alternative is that the observed allele simply evolved independently within *A. tuberculatus*.

Screening of progeny derived from *A. tuberculatus* plants did not result in the identification of hybrids. Similar levels of fecundity were reported in crosses between *A. tuberculatus* and *A. palmeri*, with low hybrid frequency (Franssen et al. 2001). Furthermore, many seeds produced by Franssen and others were attributed to the rare ‘partially monoecious’ plants observed within their experiment, hereafter referred to as ‘autopollination’ (Franssen et al. 2001). In *A. tuberculatus*, gender is determined by a single loci, where males are heterogametic (Montgomery et al. 2019). Therefore, pollen contamination is expected to result in a 1:1 ratio of male to female progeny, whereas seed derived from autopollination would result in completely female progeny. Indeed, we observed a skew towards female progeny when *A. tuberculatus* was allowed to cross only with *A. albus* (Table 3). We suspect that male progeny were obtained as a result of pollen contamination.

Apomixis also has been suggested to explain seed production observed in isolated *A. palmeri* females (Ribeiro et al. 2014). Indeed, apomixis would produce the same gender ratio (all females) as expected due to autopollination. Dioecy can be viewed as a limiting factor for the colonization of a new region. As *A. tuberculatus* is noted as an exceptional colonizer, a mechanism to overcome the limitations of dioecy may not be unexpected. However, the mechanism through which seed production is mediated is impactful for a developing population. Genetic segregation would be expected under the autopollination hypothesis, which would result in diverse progeny. In contrast, each progeny produced through the apomixis hypothesis would be genetically identical. A mechanism that promotes genetic segregation may be advantageous towards adaptation to new environments. Indeed, testing these hypotheses is straightforward: a heterozygous marker under isolation should not segregate under apomixis, but should segregate under autopollination. Preliminary attempts to identify a heterozygous loci within the selected parent plants of this study were not successful.

There are benefits and costs associated with interspecific hybridization (Chunco 2014). An invading species can rapidly gain access to adaptation traits to the new habitat, or the mixture produced could be less fit than either parent. As the range of *A. tuberculatus* continues to expand, fundamental questions remain. Is this expansion, and subsequent displacement of native species, mediated wholly through the genetic variation within the species, or obtained from outside gene flow between species? While molecular surveys provide insight into these questions, the hypotheses must then be experimentally examined. Here, we conclude that hybridization between *A. tuberculatus* and *A. albus* can happen, though at a notably low frequency in comparison to other interspecific crosses within the genus. Furthermore, due to sterility or near-sterility observed in both the initial hybrids and their progeny, widescale introgression of *A. albus* into *A. tuberculatus* seems unlikely in naturalized conditions. Theoretically, however, herbicide regimes within production agriculture could generate a sufficient selection pressure for novel herbicide-resistance traits, driving gene introgression between the species.

## Acknowledgements

We thank Dr. A Lane Rayburn for help with flow cytometry.

## Declarations

### Ethics approval

Not applicable.

### Consent to participate

Not applicable.

### Consent for publication

All authors consent to this research being published.

### Availability of data and material

Datasets and material used or generated during the current study are available from the corresponding author on reasonable request.

### Code availability

Not applicable.

### Authors’ contributions

Patrick J Tranel conceived the study. Material preparation and data collection were performed by Laura A Chatham, Danielle M McCormick, and Brent P Murphy. The first draft of the manuscript was written by Brent P Murphy. All authors read and approved the final manuscript.

## Online appendix items

**Supplementary Figure 1:**
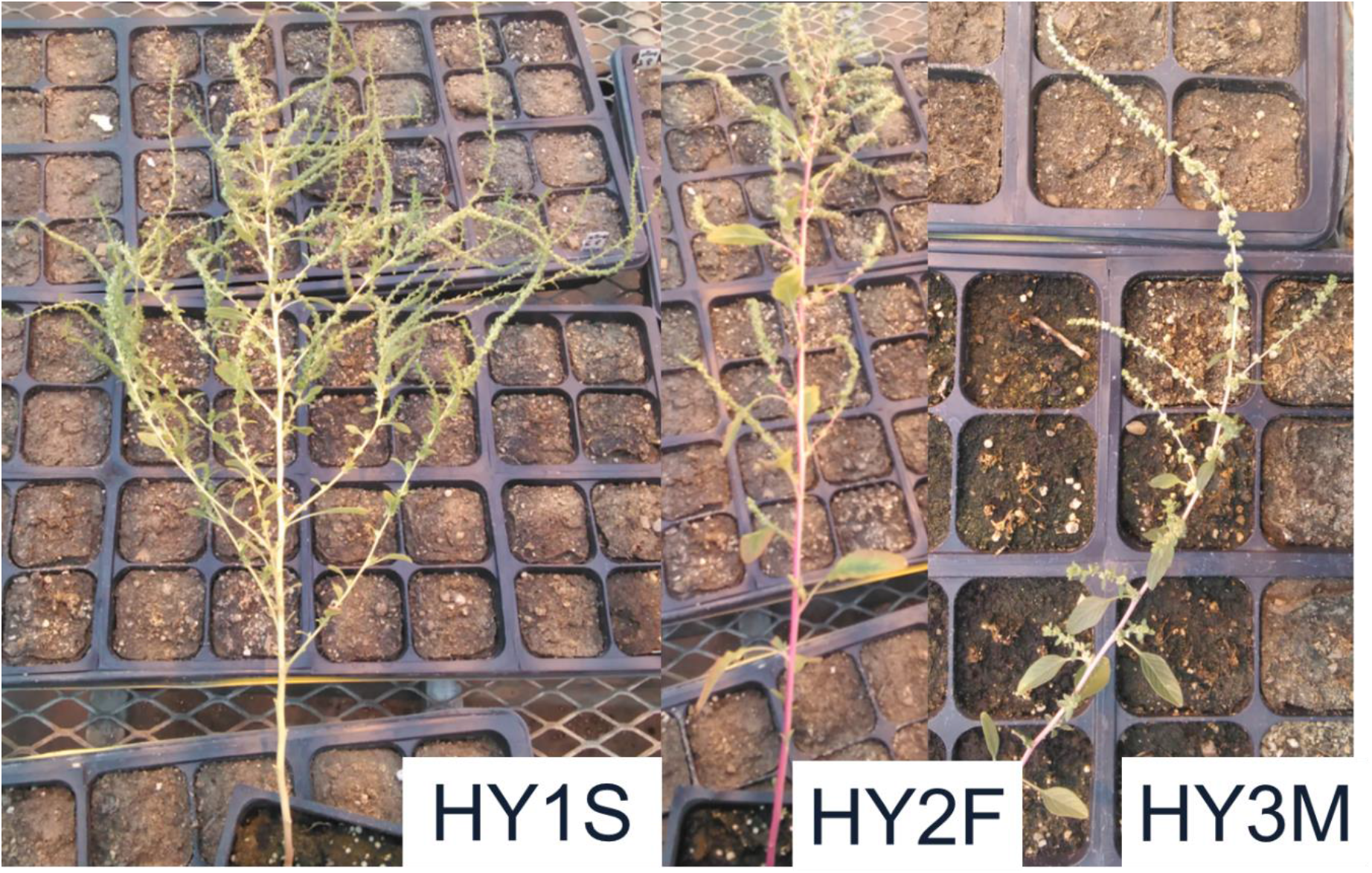
Morphology of *Amaranthus. albus* x *Amaranthus tuberculatus* hybrids.

**Supplementary Figure 2:**
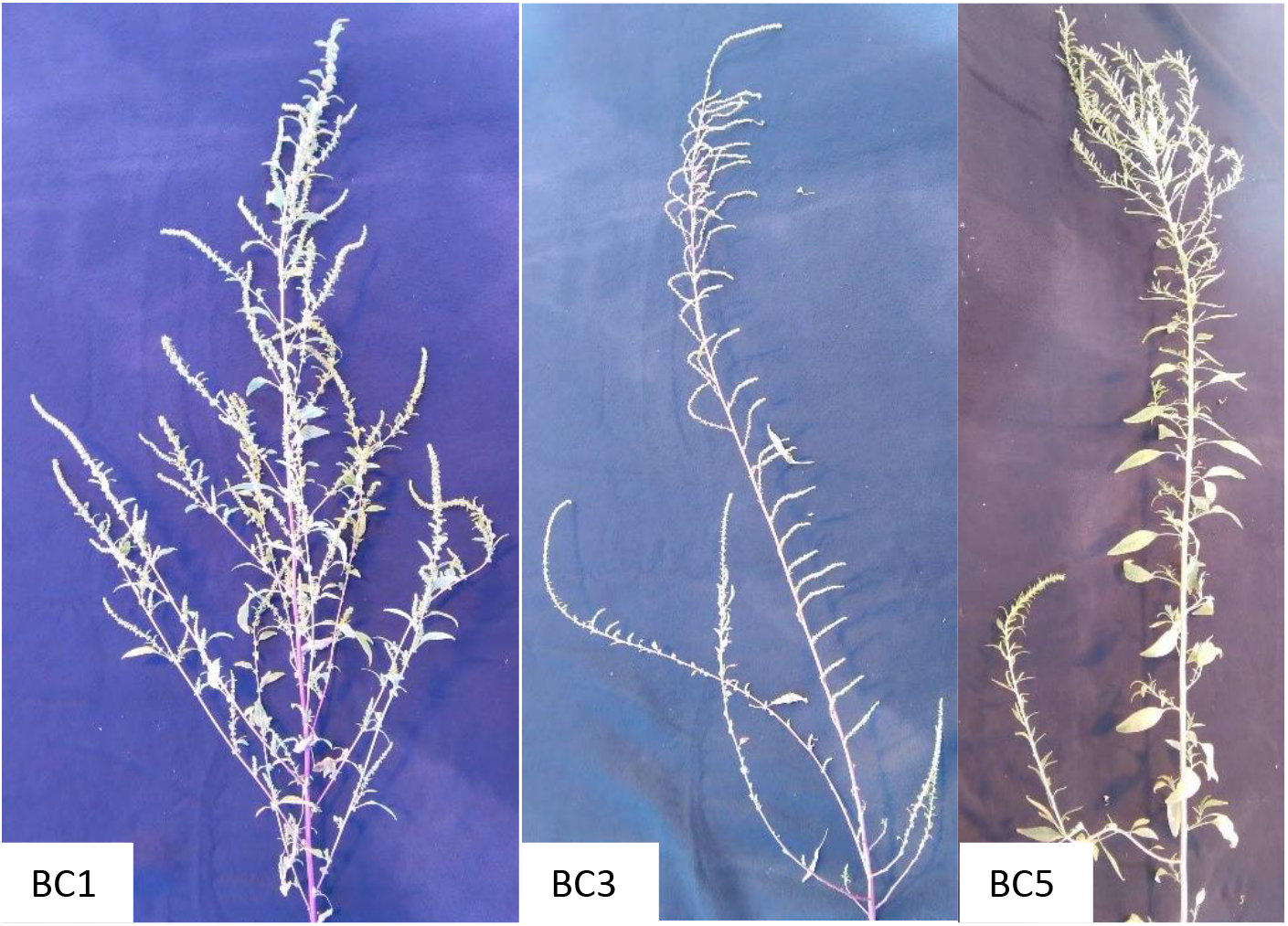
Morphology of backcross [HY2F (see Supplementary Figure 1) x *Amaranthus tuberculatus*] progeny. A, HY1; B, HY3; C, HY5.

